# Coalescence and linkage disequilibrium in facultatively sexual diploids

**DOI:** 10.1101/232405

**Authors:** Matthew Hartfield, Stephen I. Wright, Aneil F. Agrawal

**Author notes:** Website for.simulation code:* http://github.com/MattHartfield/FacSexCoalescent.

## Abstract

Under neutrality, linkage disequilibrium (LD) results from physically linked sites having non-independent coalescent histories. In obligately sexual organisms, meiotic recombination is the dominant force separating linked variants from one another, and thus in determining the decay of LD with physical distance. In facultatively sexual diploid organisms that principally reproduce clonally, mechanisms of mitotic exchange are expected to become relatively more important in shaping LD. Here we outline mathematical and computational models of a facultative-sex coalescent process that includes meiotic and mitotic recombination, via both crossovers and gene conversion, to determine how LD is affected with facultative sex. We demonstrate that the degree to which LD is broken down by meiotic recombination simply scales with the probability of sex if it is sufficiently high (much greater than 1/*N* for *N* the population size). However, with very rare sex (occurring with frequency on the order of 1/*N*), mitotic gene conversion plays a particularly important and complicated role because it both breaks down associations between sites and removes within-individual diversity. Strong population structure under rare sex leads to lower average LD values than in panmictic populations, due to the influence of low-frequency polymorphisms created by allelic sequence divergence acting in individual subpopulations. These analyses provide information on how to interpret observed LD patterns in facultative sexuals, and determine what genomic forces are likely to shape them.

## Introduction

Coalescent theory is a powerful mathematical framework that is used to determine how natural selection and demographic history affect genetic diversity (Kingman 1982; Rosenberg and Nordborg 2002; Hein *et al*. 2005; Wakeley 2009). Traditional coalescent models assume that the population is obligately sexual, but there has been less attention on creating models that account for different reproductive modes. While the coalescent with self-fertilisation has been extensively studied (Nordborg and Donnelly 1997; Nordborg 1997, 2000; Nordborg and Krone 2002), little theory exists on coalescent histories in organisms with other mixed reproductive systems.

Previous theory has investigated genetic diversity in facultatively sexual diploid organisms, which reproduce via a mixture of sexual and parthenogenetic reproduction (Brookfield 1992; Burt *et al*. 1996; Balloux *et al*. 2003; Bengtsson 2003; Ceplitis 2003). A general result arising from this work is that when an organism exhibits very rare population-level rates of sex 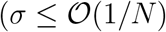, for *σ* the probability of sex and *N* the population size), they will exhibit ‘allelic sequence divergence’ where both alleles within a diploid individual accumulate distinct polymorphisms from each other (Mark Welch and Meselson 2000; Butlin 2002). Hartfield *et al*. (2016) subsequently investigated a coalescent model of facultative sexuals, and quantified how the presence of gene conversion can reduce within-individual diversity to less than that expected in sexual organisms, contrary to the effects of allelic sequence divergence. Hence these results provide a potential explanation as to why allelic divergence is not widely observed in empirical studies of facultatively sexual organisms (reviewed in Hartfield (2016)).

However, this analysis only modelled the genetic history at a single, non-recombining locus. Here, genealogies only greatly differed from those in obligately sexual organisms at very low frequencies of sex 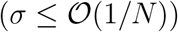. As a consequence, methods to estimate the frequency of sex can only do so based on the degree of allelic sequence divergence, and are expected to be ineffective if the frequency of sex is greater than 1/*N* and/or gene conversion is prevalent (Ceplitis 2003; Hart-field *et al*. 2016). In contrast, many facultatively sexual organisms exhibit much higher occurrences of sex. Pea aphids reproduce sexually about once every ten to twenty generations (Jaquiéry *et al*. 2012), while Daphnia undergo one sexual generation and five to twenty asexual generations a year (Haag *et al*. 2009). The wild yeast *Saccharomyces paradoxus* has an outcrossing frequency of 0.001; while low, this value is four orders of magnitude higher than 1/*N_e_* (Tsai *et al*. 2008). If we wish to create a general coalescent model that can be used to estimate rates of sexual reproduction in species undergoing more frequent sex, then we need to increase the power of this coalescent process to consider how patterns of genetic diversity at multiple loci are affected with facultative sex.

This is achievable by considering how genealogies of multiple sites correlate along a chromosome. Two completely linked sites will reach a common ancestor in the past at the same time, so will share the same gene genealogy. However, if a recombination event (e.g., via meiotic crossing over) were to separate the sites, each sub-segment may have different genetic histories (Hudson 1983). Breaking apart correlations between sites is reflected with lower linkage disequilibrium, which can be measured from genomic data (Griffiths 1981; Hudson and Kaplan 1985; Hudson 1990; Simonsen and Churchill 1997; McVean 2002). Gene conversion can also break apart correlations between sites through transferring genetic material across DNA strands (Wiuf and Hein 2000).

As meiotic crossing over occurs during sexual reproduction, one may expect that the extent to which linkage disequilibrium is broken down should scale with the probability of sex (see Nordborg (2000) for a related argument for the coalescent with self-fertilisation). Tsai *et al*. (2008) used this logic to calculate the frequency of sex in the yeast *Saccharomyces paradoxus*, by comparing effective population sizes inferred from linkage disequilibrium decay (which should scale with the meiotic recombination rate, and therefore the rate of sex) with those from nucleotide variation (which should be independent of sex if sufficiently high). Lynch *et al*. (2017) used similar arguments to conclude that even though the facultatively sexual water flea *Daphnia pulex* has a lower overall crossover recombination rate than *Drosophila melanogaster*, it has a higher crossover rate when sex does occur.

However, the logic used in these studies assumes that the frequency of sex only affects occurrences of meiotic crossing over. Low rates of sex also distort the underlying genealogies, leading to subsequent events (including allelic sequence divergence or removal of diversity via gene conversion) that also affect how polymorphisms are correlated along haplotypes. Hence these approaches may become problematic in species exhibiting low rates of sexual reproduction, or if gene conversion is an important force in shaping genetic diversity, as observed in empirical studies of facultative sexuals (Crease and Lynch 1991; Schön *et al*. 1998; Normark 1999; Schön and Martens 2003; Flot *et al*. 2013).

We describe both mathematical theory and a routine for simulating multi-site genealogies with facultative sex, allowing for both meiotic and mitotic crossover recombination and gene conversion. We use these new models to investigate how linkage disequilibrium patterns are affected in facultatively sexual organisms, and how these results can be used to infer rates of sex from genome data. Specifically, we investigate when the breakdown of linkage disequilibrium scales with sex, as predicted by intuition, and when this logic does not hold.

## Overview of the facultative-sex coalescent and re-combination events

Our primary goal is to examine how different frequencies of sex affects linkage disequilibrium. Heuristically speaking, the expected strength of disequilibrium depends on the probability that two sampled haplotypes (hereafter ‘samples’) co-alesce before either haplotype is disrupted by recombination. Before presenting the formal model, we begin by discussing how facultative sex affects coalescence and then recombination.

In the standard coalescent, each member of a set of (non-recombining) samples can be thought of as travelling independently backward in time through the generations. A coalescence event occurs if two samples independently “choose” the same parental allele as their ancestor. The waiting time until the next coalescence depends only the number of remaining samples but, importantly, not on “where” the samples are currently found (i.e., in which individual organisms). However, for diploids with a low frequency of sex, the “where” information is crucial (Bengtsson 2003; Ceplitis 2003; Hartfield *et al*. 2016). For example, two samples can be the two haplotypes found in a single diploid individual (which we denote as ‘a paired sample’) or they can each come from different individuals (which we denote as ‘two unpaired samples’). The two haplotypes within a paired sample do not travel back in time independently, rather they travel together for as long as reproduction is asexual. Coalescence between them is not possible for all the asexual generations they remain paired (ignoring gene conversion). A sexual event splits a paired sample into two unpaired samples that can then coalesce in a subsequent generation. For this reason, paired samples are expected to have longer average coalescence times than unpaired samples and low sex increases average coalescence time compared to high sex (Bengtsson 2003; Ceplitis 2003; Balloux *et al*. 2003). However, if the frequency of mitotic gene conversion is high relative to the frequency of sex, then these predictions are reversed (Hartfield *et al*. 2016). In this case, paired samples can coalesce faster than unpaired samples because each generation the samples are paired provides an opportunity for coalescence via mitotic gene conversion.

In sum, low sex in diploids requires a ‘structured’ coalescent approach because paired and unpaired samples behave differently; this structure affects the distribution of coalescence times (including the mean and variance) and is sensitive to the amount of mitotic gene conversion. Technically, this structure occurs even in diploids that are obligately sexual; however, the coalescent can be safely modelled ignoring the structure because the time spent in “paired” states is infinitesimally brief on the coalescent timescale when sex is common. To affect coalescence times, sex must be sufficiently uncommon, i.e. 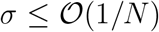, where *N* is the population size and *σ* is the fraction of offspring sexually produced each generation via the random union of gametes (i.e., *σ* = 1 represents obligate sex and *σ* = 0 obligate asexuality; see Table 1 for a list of symbol definitions).

**Table 1:** Glossary of Notation.

| Symbol | Usage |
| --- | --- |
| $N$ | Diploid population size (with $2N$ haplotypes), denoted $N_T$ if measured across a subdivided population |
| $f_A, f_B$ | Frequency of derived allele at site $A, B$ |
| $f_{AB}$ | Frequency of haplotypes carrying derived allele at both sites |
| $D_{AB}$ | Disequilibrium between two sites, $f_{AB} - f_A f_B$ |
| $r^2$ | Standardised linkage disequilibrium, $D_{AB}/(f_A(1 - f_A)f_B(1 - f_B))$ |
| $r_d^2$ | ‘Ratio of means’ measure of linkage disequilibrium, $E[D_{AB}]/(E[f_A(1 - f_A)f_B(1 - f_B)])$ |
| $t_A(ij)$ | Coalescent time at site $A$ (if sampled from haplotypes $i, j$) |
| $\sigma$ | Fraction of offspring produced via sex |
| $c, c_A$ | Probability of meiotic (mitotic) crossing over between two sites |
| $\tilde{c}, \tilde{c}_A$ | Probability of meiotic (mitotic) crossing over between two adjacent sites |
| $\gamma_1, \gamma_2$ | Probability of mitotic gene conversion covering one, two sites (analytical model) |
| $\gamma_{1S}, \gamma_{2S}$ | Probability of meiotic gene conversion covering one, two sites (analytical model) |
| $g, g_S$ | Probability of mitotic (meiotic) gene conversion initiating at a site |
| $\Omega$ | Population-level frequency of sex, $2N\sigma$ |
| $\rho, \rho_A$ | Population-level rate of meiotic (mitotic) crossing over, $4Nc$ ($4Nc_A$) |
| $\tilde{\rho}_A$ | Population-level rate of mitotic crossing over between two adjacent sites, $4N\tilde{c}_A$ |
| $\Gamma_1, \Gamma_{1S}$ | Population-level rate of mitotic (meiotic) gene conversion affecting a single site, $4N\gamma_1$ ($4N\gamma_{1S}$) |
| $G$ | Population-level rate of gene conversion initiation, $4Ng$ |
| $\lambda, \lambda_S$ | Average length of mitotic (meiotic) gene conversion event |
| $L$ | Number of sampled sites; $L - 1$ is number of breakpoints |
| $Q, Q_S$ | Number of breakpoints in units of average gene conversion length (e.g. $Q = (L - 1)/\lambda$ for mitotic gene conversion) |
| $R$ | Population-level meiotic crossing rate (simulation), $4N\tilde{c}(L - 1)$ |
| $\Gamma$ | Population-level mitotic gene conversion rate (simulation), $4Ng(L - 1)$ |
| $\phi$ | Ratio of sex to mitotic gene conversion acting at a single site, $(\Omega Q)/\Gamma$ |
| $\mu$ | Mutation rate over $L$ sites |
| $\theta$ | Population level mutation rate, $4N\mu$ |
| $m$ | Probability of migration (island model) |
| $M$ | Population-level rate of migration, $2N_T m$ |
| $p_1, p_2$ | Probability of one, two gene conversion breakpoints within sample |

Different sites along a genetic segment can have different genealogical histories as long as there is some recombination. Low sex affects recombination in several ways. We consider both crossing-over (the reciprocal exchange of genetic material between two haplotypes) and gene conversion (where genetic material is copied from one haplotype to its homolog). When sexual reproduction is rare, the frequency of meiotic recombination will necessarily be low. Mitotic crossovers and mitotic gene conversion can then become important for two reasons. First, in comparison to meiotic recombination, mitotic recombination becomes a relatively more important route of genetic exchange as meiosis becomes rare. Second, in paired samples (which are only an important consideration when sex is low), mitotic recombination can either lead to gene exchange (the splitting of a multisite sample into separate pieces) or coalescence.

Figure 1 outlines the possible outcomes for recombination under facultative sex. Going back in time, sex involving a meiotic crossover will transform an unpaired sample into a paired sample (i.e., the unpaired sample descended from the two homologs in the parent; Figure 1(a)). For a paired sample, sex segregates the two samples into separate parents, creating two unpaired samples (Figure 1(b)). However, if a crossover also occurred on one of these samples, then the affected sample becomes a paired sample in the parent; the overall outcome is a new paired sample in one parent (each containing a section of ancestral material), and one unpaired sample in the other parent (Figure 1(c)). Mitotic crossovers can also act in paired samples unaffected by sex, swapping genetic material between homologues (Figure 1(d)).

**Figure 1:**
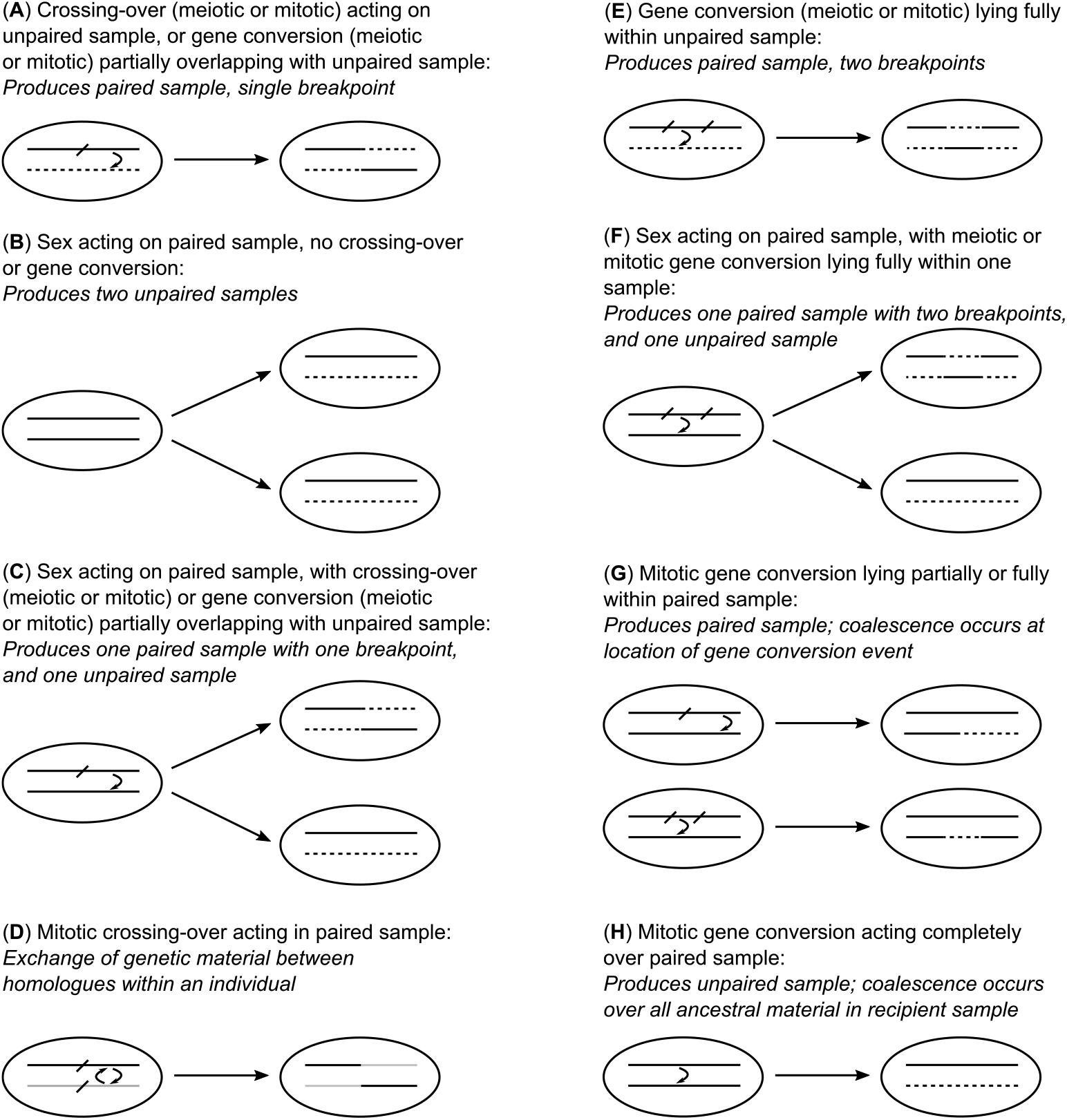
Schematic of different outcomes following gene exchange in the facultative-sex coalescent. Solid lines represent ancestral material; dotted lines represent non-ancestral material. Outcomes are described for (a) meiotic crossing over acting on an unpaired sample, or gene conversion acting on an unpaired sample that only partially overlaps with the haplotype; (b) sex acting on a paired sample with no crossing over or gene conversion; (c) both sex and either crossing over, or gene conversion that only partially overlaps with the haplotype, acting on a paired sample; (d) mitotic crossing over acting on a paired sample; (e) gene conversion (meiotic or mitotic) acting on an unpaired sample, which fully lies within the sampled haplotype; (f) both sex and gene conversion (lying fully within a haplotype) acting on a paired sample; (g) mitotic gene conversion acting on a segment of a paired sample; (h) mitotic gene conversion acting over the entire length of a paired sample (or over all remaining extant material).

Gene conversion can affect a sample in several ways, where (i) gene conversion initiates outside a tract of ancestral material but finishes within it; (ii) gene conversion begins within a tract of ancestral material but extends beyond it; (iii) both conversion breakpoints lie within ancestral material; or (iv) gene conversion acts over all ancestral material in a paired sample (see Wiuf and Hein (2000) for a detailed discussion of the coalescent with gene conversion applicable to obligately sexual diploids). If gene conversion acts on an unpaired sample, then it becomes a paired sample with each haplotype carrying a segment of ancestral material, which is a similar outcome to that following a crossover (Figure 1(e)). There are either one or two breakpoints, depending on whether gene conversion lies partly or fully within ancestral material. If acting on paired samples, the outcome depends on whether sex has segregated the samples into different individuals. If so then one of the two parents contains a paired sample with each part carrying ancestral material (Figure 1(f)). If not, then a segment of one sample coalesces into the other (Figure 1(g)). Finally, mitotic gene conversion acting completely over a paired sample reproducing asexually causes complete coalescence of one paired sample, converting it into an unpaired sample. This outcome is equivalent to ‘gene conversion’ for the single-site coalescent model (Hartfield *et al*. 2016) (Figure 1(h)).

Overall, facultative sex will affect linkage disequilibrium for at least three reasons. First, the population-level rate of meiotic recombination will be proportional to the frequency of sexual reproduction. Second, when sex becomes very rare, the rate and patterns of coalescence change substantially, which is important because disequilibrium is affected by the rate of recombination relative to coalescence. Third, in the low-sex regime, mitotic gene conversion can become important as it becomes a key coalescence mechanism for a paired sample; alternatively, a single haplotype within an individual can be separated (with either one or two break-points) via gene conversion.

## Two-site analytical model

A commonly used metric of linkage disequilibrium is (Hill and Robertson 1968):

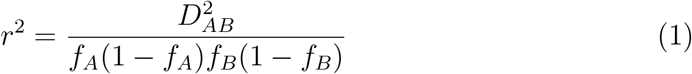

where *f_i_* is the frequency of the derived allele at site *i* (*i* = *A* or *B*), and *D_AB_* = *f_AB_* − *f_A_f_B_* with *f_AB_* being the frequency of haplotypes carrying the derived allele at both sites. A related quantity

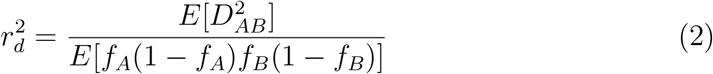

has been studied in analytical neutral models (Ohta and Kimura 1971; Weir and Hill 1986; McVean 2002) (we use 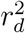 to represent this quantity, rather than the traditional symbol 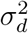, to avoid confusion with *σ* that is used to parameterize the frequency of sex in this analysis). 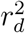 overestimates the expected value of *r*^2^ but the discrepancy is reduced if it is only applied to sites where the minor allele is not too rare (McVean 2002). In the classic analyses, which is applicable to obligately sexual diploids:

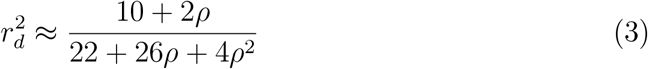

where *ρ* = 4Nc with *c* being the per-generation probability of meiotic crossing over between two sites. McVean (2002) showed that a coalescence approach can be used to derive this result, demonstrating that 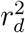 is a function of the covariance in coalescence times between two sites. The goal here is to quantify how 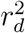 is altered by facultative sex. We use the coalescent approach of McVean (2002) for a two-site model in a diploid population of size *N*. Two samples at each of two sites in a diploid model can occur in 17 different states, as outlined in Figure 2. In the traditional haploid model, only 7 states are necessary, but here we must consider the full 17-state model to consider the pairing of haplotypes. The model is presented in detail in Section A of Supplementary Mathematica File S1, with an overview provided in Figure 2.

**Figure 2:**
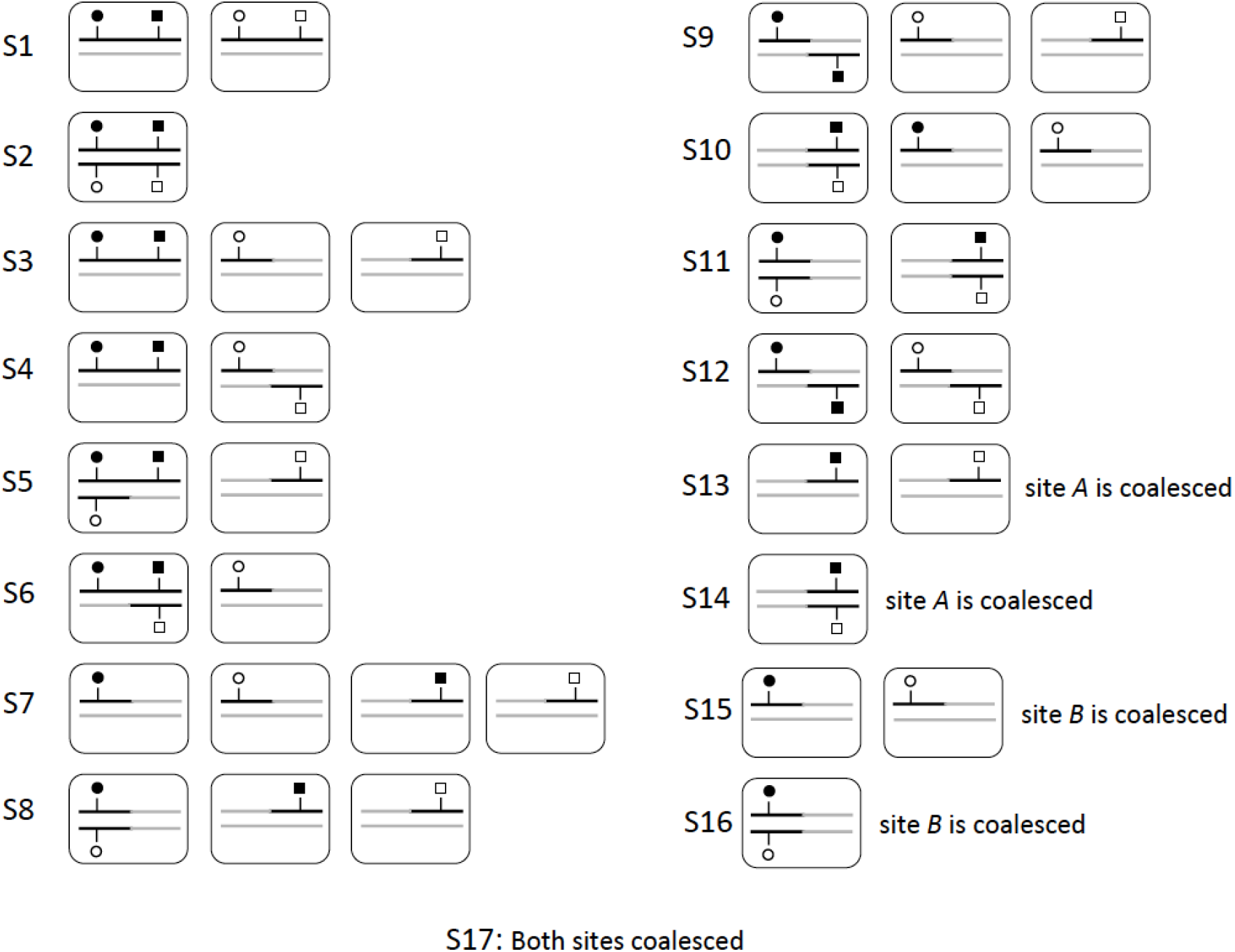
The 17 possible states for two copies of each of two sites, for the analytical model. Each rounded rectangle represents a separate diploid individual. The two focal copies of the *A* site are represented by open and closed circles. The two focal copies of the *B* site are represented by open and closed squares. The shading of symbols, i.e., open vs. closed, has no meaning other than to distinguish focal copies. Haplotypes or parts of haplotypes that do not carry ancestral material (i.e., not carrying focal copies) are shown in grey. Coalesced sites are not shown.

The first key step in constructing the model is to derive the transition matrix giving the probabilities (going backwards in time) of changing states. These probabilities depend on the biology of reproduction and inheritance. If meiosis occurs, there is a crossover between sites *A* and *B* with probability *c*. The probability of a mitotic crossover is *c_A_* per generation (which does not require meiosis). Regardless of reproductive mode, mitotic gene conversion can occur. With probability *γ*_2_ there is a mitotic gene conversion event whose tract length covers both sites. With probability *γ*_1_ a mitotic gene conversion event occurs where one end of the gene conversion tract lies at the breakpoint between the two sites, and the other end lies beyond them (*γ*_1*S*_ is the analogous probability for a meiotic gene conversion event, conditional on meiosis occurring). It is worth noting that *γ*_1_ enters the transition matrix for two separate reasons. It determines the probability that one site coalesces via mitotic gene conversion (e.g., transition from state S2 to S14; see Figure 2) and it determines the probability that samples at different sites on the same haplotype get split onto separate haplotypes by mitotic gene conversion (e.g., transition from state S3 to S9; see Figure 2). Note that gene conversion involving one site is functionally equivalent to a crossing-over event. Using these parameters, the construction of the transition matrix is tedious but straightforward. In contrast, *γ*_2_ only enters transitions involving coalescence affecting one or both sites. Using first-step analysis (Wakeley 2009, Chapter 7) and following McVean (2002), we construct a system of equations for the expected value of the product of coalescent times at the two sites, given their current state *z*. These equations capture the expected time for the system to move out of the state *z*, before calculating the expected coalescent time of either one or both sites, given the new state *k*. These equations have the form:

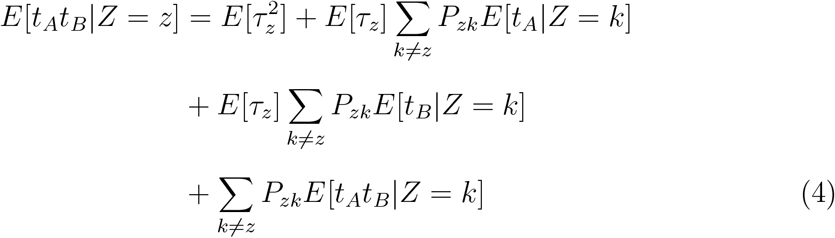

where *τ_z_* is the time to exit state *z, P_zk_* is the probability that the system moves from state *z* to state *k* conditional on leaving state *z*, and *E*[*t_x_*|*Z* = *k*] is the expected time to coalescence of site *x* given it is currently in state *k*. As described in Section A of Supplementary Mathematica File S1, these components can be calculated from the transition matrix, either directly for discrete time or after appropriate transformation for the continuous time approximation (Möhle 1998; Wakeley 2009).

Following McVean (2002):

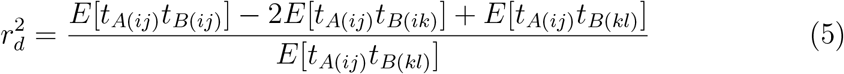

where *E*[*t_A(ij)_t_B(kl)_*] is the expected product of the coalescent times at site *A*, where the two copies are sampled from haplotypes *i* and *j*, and at site *B*, where the two copies are sampled from haplotypes *k* and *l* (and different indexes denote other haplotype samples). Analogous to measuring 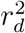 from haploids where each haplotype represents an independent sample, we calculate 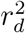 assuming that each haplotype comes from a different individual (Figure 2) so that the three terms in the numerator represent coalescence times from states S1, S3, and S7.

We first consider the case of partial asexuality where sex may be rare at the individual level but is not too rare at the population level (i.e., 0 < *σ* ≤ 1 but *Nσ* ≫ 1). We find

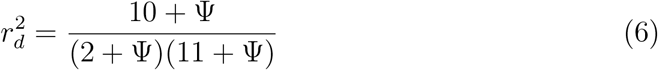

where Ψ = *ρσ* + *ρ_A_* + (1/2)Γ_1_ + (1/2)*σ*Γ_1*S*_ with scaled parameters *ρ* = 4*Nc*, *ρ_A_* = 4*Nc_A_*, Γ_1_ = 4*N*_*γ*1_, and Γ_1*S*_ = 4*N*_*γ*1*S*_. Simplifying the model by ignoring gene conversion and mitotic crossing over (Γ_1_ = Γ_1*S*_ = *ρ_A_* = 0), the result above is the same as the obligate sex result (Equation 3) but using an effective scaled crossover rate *ρσ* in place of *ρ* (Figure 3(a)).

**Figure 3:**
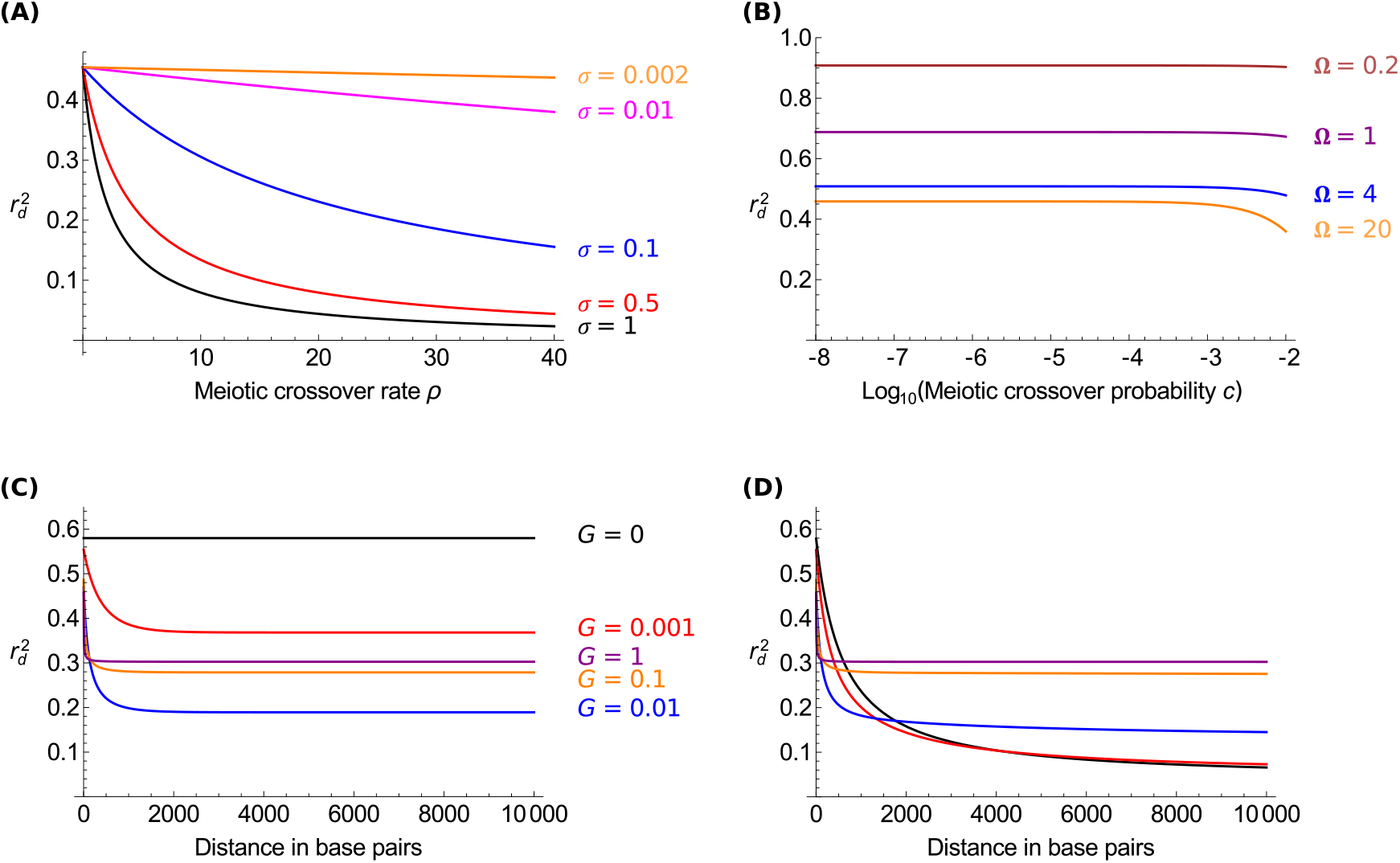
(a) Linkage disequilibrium, measured as 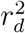, when sex is high at the population level (i.e., 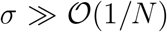), measured as a function of the meiotic crossover rate *ρ*. Different frequencies of sex (*σ*) are shown. Other parameters: *c_A_* = *γ*_1_ = *γ*_2_ = *γ*_1*S*_ = 0. (b) 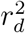 when the frequency of sex is low 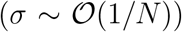 and the only haplotype disrupting force is meiotic crossing over (*c* > 0). Different rates of sex (Ω = 2*Nσ*) are shown. Other parameters: *c_A_* = *γ*_1_ = *γ*_2_ = *γ*_1*S*_ = 0. (c, d) 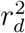 when the frequency of sex is low 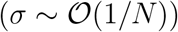 as a function of the distance between two sites (measured in basepair distance), for different levels of mitotic gene conversion (*G* = 4*N_g_*). Results are shown without (c) and with (d) mitotic crossing-over (i.e., 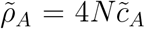 with 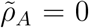 in (c) and 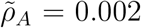 in (d)). Other parameters: λ = 500; Ω = 2 (see Figure A in Supplementary File S2 for similar plots using different Ω values).

We next consider the case where sex is rare at the population level, 2*Nσ* → Ω as *N* → ∞. In the absence of mitotic gene conversion or mitotic crossing over (Γ_1_ = *ρ_A_* = 0) then:

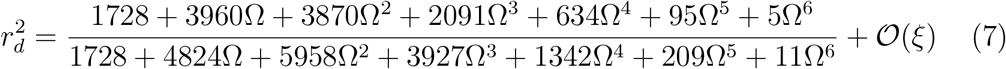

where the rates of disruptive meiotic processes *c*, 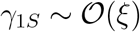 with being a small term (|*ξ*| ≪ 1). Equations 6 and 7 differ in several important ways. First their maximum values, and the conditions to achieve these maxima, differ. The maximum value of Equation 6 occurs as the haplotype disrupting forces (crossovers and gene conversion) become small, i.e., 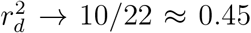 as Ψ → 0. In contrast, the maximum value of Equation 7 occurs as sex becomes increasingly rare, i.e., 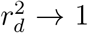 as Ω → 0. Second, 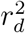 in Equation 6 has a strong dependence on physical distance because disruption via crossover or gene conversion is an increasing function of distance, i.e., Ψ is implicitly an increasing function of distance. In contrast Equation 7 is very weakly dependent on physical distance through terms of 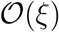 (Figure 3(b)).

Equation 7 assumes no mitotic gene conversion or mitotic crossing over but important changes to 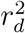 occur with either of these processes. An analytical approximation can be obtained but the expression is unwieldy (Section A of Supplementary Mathematica File S1). Both types of mitotic gene conversion events, represented via Γ_1_ = 4*N*_*γ*1_ and Γ_2_ = 4*N*_*γ*2_, as well as mitotic crossing over (*ρ_A_* = 4*N_c_A__*) affect the leading-order term for 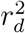 and are functions of the distance between sites. Mitotic crossing over can be modelled as a linear function of distance 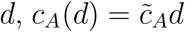. Using a standard assumption of exponentially distributed gene conversion tract lengths (Wiuf and Hein 2000), the probabilities of mitotic gene conversion are given by *γ*_1_(*d*) = 2*g*λ(1 − exp(−*d*/λ)) and *γ*_2_(*d*) = *g*λexp(−*d*/λ) where λ is the average tract length and *g* is the probability of gene conversion initiation per base pair (more precisely, per breakpoint between adjacent base pairs). The derivation is provided in Section A of Supplementary Mathematica File S1.

Figure 3(c) shows that 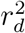 declines with physical distance when there is mitotic gene conversion but no mitotic crossing over. Note that 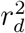 does not decline down to 0 with distance as it does in the classic model (Equation 3) of meiotic crossing over. Because gene conversion probabilities change slowly for *d*/λ > 2, there is little decline in 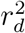 beyond this point. Surprisingly, 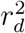 is not always a monotonically declining function of the probability of gene conversion initiation, *g* (or the scaled parameter *G* = 4*N_g_*), especially when *d* > λ (Figure B(a) in Supplementary File S2). Consequently, a species with a lower frequency of gene conversion events (i.e., smaller *g*) can have larger 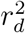 for small *d* but smaller 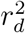 for large *d* compared to an otherwise similar species with larger *g* (Figure 3(c)). This behaviour of 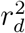 with respect to *g* is likely due to the dual (and conflicting) roles of gene conversion in increasing both the probability of coalescence and disruption of haplotypes. In contrast, mitotic crossing over, which only affects haplotype disruption, affects 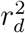 monotonically as expected (Figure B(b) in Supplementary File S2). The addition of mitotic crossing over reduces the minimum value of 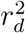 (Figure 3(d)). Even with mitotic crossing over, 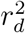 does not go to zero at large distances and can be considerably greater than zero when gene conversion is high (see Section A of Supplementary Mathematica File S1). The minimum value reached by 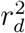 is independent of the rate of mitotic crossing over (provided it is not zero) though the distance at which the minimum is reached is shorter with higher rates of mitotic crossing over.

## Simulation Algorithm

We have previously developed an algorithm to build genealogies of facultative sexual organisms at a single non-recombining locus (Hartfield *et al*. 2016). This algorithm simulates genealogies of *n* samples, of which 2x are paired and the remaining *y* = *n* − 2*x* samples are unpaired. The algorithm proceeds in a similar manner as other coalescent simulations, in that it tracks the genetic histories of samples into the past, sequentially enacting events that affect the genetic history (e.g. coalescence, sexual reproduction). The relative probability of each event occurring per generation is used to determine what the next event is, and at which time in the past it arises. To further investigate the effects of facultative sex on linkage disequilibrium, we extended this previous routine to consider coalescent histories of multiple sites, and how various recombination phenomena affect how genetic histories are correlated along chromosomes. In Appendix A we describe how the crossover routine of Hudson (1983) and the gene conversion routine of Wiuf and Hein (2000) are extended to consider the effects of facultative sex. As a consequence, the updated coalescent simulation now models the effects of meiotic and mitotic recombination on facultative sex, the outcomes of which are summarised in Figure 1.

### Measuring linkage disequilibrium from simulations

We used the updated coalescent simulation to calculate expected linkage disequilibrium in facultatively sexual organisms. Following a single simulation of a coalescent process, a series of *j* genealogies are produced, one for each non-recombined part of the genetic segment. Polymorphisms are added to each branch of the genealogy, drawn from a Poisson distribution with mean (1/2)*θ_j_τ_i,j_*, for *θ_j_* = 4*N_T_μ*(*l_j_/L*) the mutation rate of the segment covering *l_j_* of *L* total sites given *μ* is the mutation rate for a segment of *L* sites, and *τ_i,j_* the length of branch *i* in segment *j*.

For each simulation, we measured linkage disequilibrium *D* = *f_AB_* − *f_A_f_B_* over each pairwise combination of polymorphisms; this measure was then normalised to *r*^2^ = *D*^2^/(*f_A_f_B_*(1 − *f_A_*)(1 − *f_B_*)). Once *r*^2^ was measured over all simulations, values were placed into 20 equally-sized bins based on the distance between the two polymorphisms. However, the number of pairwise samples were different for each of the 20 bins. Samples in the last two bins produced noisy estimates of linkage disequilibrium, so we only reported linkage disequilibrium estimates from the first 18 bins. We randomly subsampled data in bins 1 to 18 so that they include the same number of pairwise comparisons as in the smallest bin that contained data, to standardise bin size per simulation. Mean values per bin were recorded for each simulation run. We then calculated the mean of means per bin over all 1,000 simulations, omitting points where data was not present in a bin for a simulation. 95% confidence intervals were calculated as 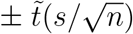 for s the standard deviation for the bin, *n* the number of points in the bin (maximum of 1,000, one for each simulation run), and 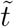 the 97.5% quantile for a *t*-distribution with *n* − 1 degrees of freedom.

### Measuring correlation in coalescence time between sites

For some cases with low sex and mitotic gene conversion, we measured the correlation in coalescence times between sites as a function of the distance between them, to investigate how these values relate to observed linkage disequilibrium patterns. For each simulation run, we obtained the number of non-recombined regions, and the times at which the ancestral segment of that region for each individual coalesced. If more than 100 segments existed, these were subsampled down to 100. We calculated the Pearson correlation in coalescent times for all segments; values were then placed into one of 20 bins based on the distance between blocks (the location of each segment was given by its midpoint). Values were only reported for the first 18 bins, with further subsampling performed on bins 1 to 18 so they contained the same number of comparisons as the smallest bin that contained data for that simulation. The mean bin value for a simulation, as well as the mean of means over all simulations and 95% confidence intervals were calculated using the same method as for linkage disequilibrium measurements.

### Data Availability

This new simulation program, *FacSexCoalescent*, along with documentation is available from *http://github.com/MattHartfield/FacSexCoalescent*. We first rebuilt the single-locus simulation program in C to greatly increase execution speed, before adding the crossover and gene conversion routines. As with the previous version of the simulation, *FacSexCoalescent* uses a timescale of 2*N* generations while `ms` uses 4*N* generations. The documentation specifies other cases where *FacSexCoalescent* inputs and outputs differ from other coalescent simulations. We performed various tests of the simulation as described in Section B of Supplementary File S2.

Supplementary File S1 is a *Mathematica* notebook of analytical derivations. Supplementary File S2 contains additional results and figures. Supplementary File S3 is a copy of the simulation code and manual. Supplemental files have also been uploaded to FigShare.

## Simulation Results

### Linkage disequilibrium with crossing over

#### High frequencies of sex

We looked at how patterns of linkage disequilibrium are affected by crossovers, when sexual reproduction is frequent (that is, the scaled rate of sex *Nσ* ≫ 1). Analytical results (Equation 6) suggest that the effect of meiotic crossovers on linkage disequilibrium is equal to that observed in an obligately sexual population with a rescaled probability *c_eff_* = *cσ* (for 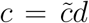 the crossover probability over distance *d*). To further investigate this pattern, we simulated genealogies over *L* = 1, 001 sites with a fixed population-level meiotic crossover rate over the entire ancestral segment 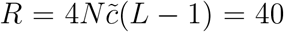, which acts during sexual reproduction. Results are reported over the first 900 sites.

Figure 4(a) plots how linkage disequilibrium decays over this region with different probabilities of sex, varying from *σ* =1 (i.e., obligate sex) to *σ* = 0.002. As expected, the decay in linkage disequilibrium is weakened with lower sex, since there exists fewer opportunities for crossovers to act (compare Figure 4(a) to analytical expectations in Figure 3(a)). We confirm that the observed decay is equivalent to an obligately sexual population with *c_eff_* = *cσ* in three ways. First, we ran equivalent (but haploid and sexual) simulations in `ms` using the rescaled crossover probability, and observed that the decay in linkage disequilibrium matches results from the facultative-sex simulation (Figure 4(b)). Second, we used the ‘pairwise’ routine in the *LDhat* software (McVean *et al*. 2002) to estimate crossover rates from facultative-sex simulation data, and observed that they scaled linearly with *σ* (Figure 4(c)). Finally, Figure 4(d) plots linkage disequilibrium values for all facultative-sex coalescent simulations as a function of the effective recombination rate, alongside the analytical expectation for 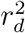 (Equation 6). *r*^2^ is calculated after removing sites with minor allele frequency < 10%, as 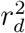 is known to overestimate *r*^2^ if all allele frequencies are considered. We see that the decay in linkage disequilibrium over all simulations is close to, but slightly overshoots the theoretical expectation (Equation 6). Similar behaviour was observed by McVean (2002, Figure 3; compare solid line to square points).

**Figure 4:**
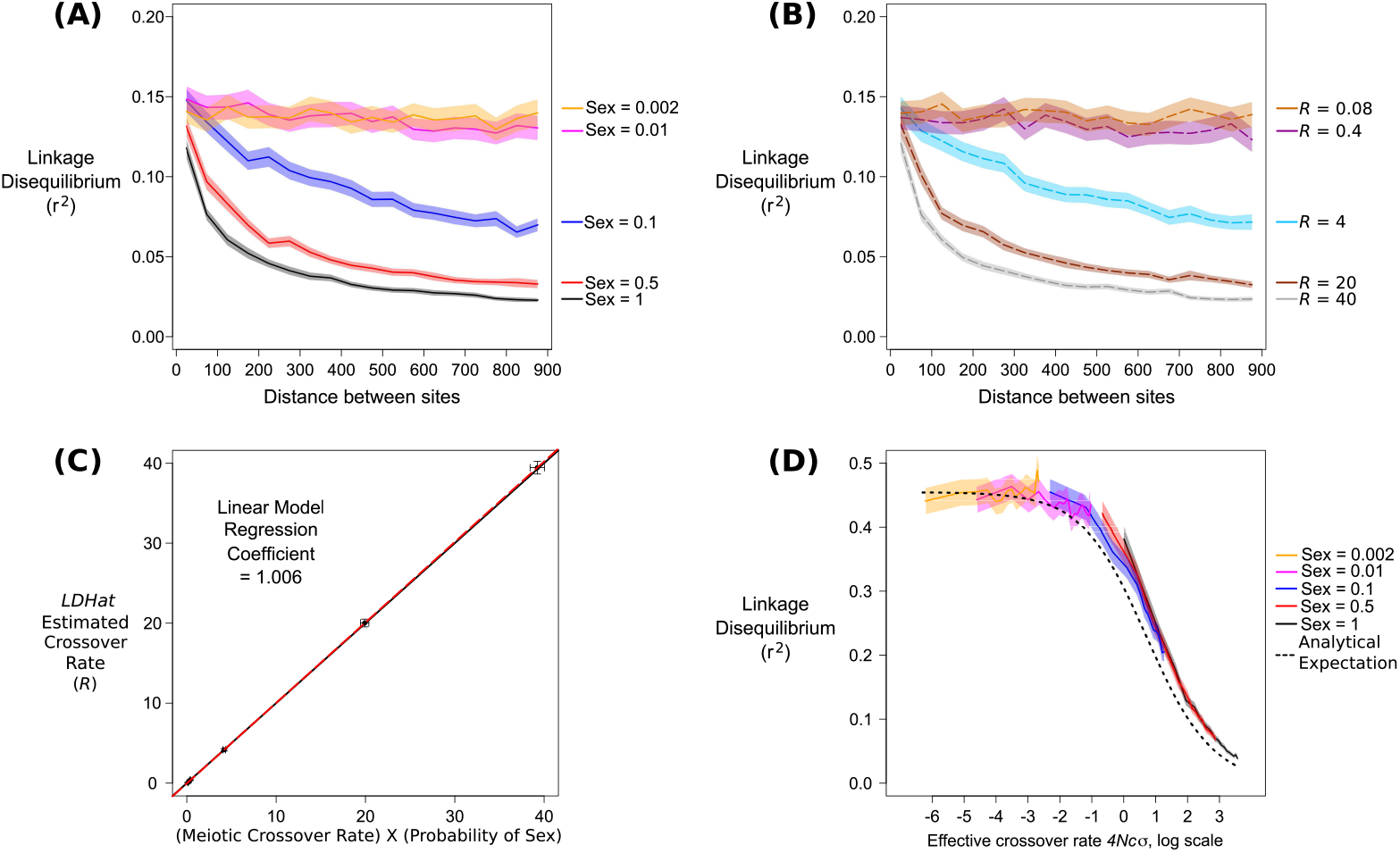
Effects of facultative, but not very low rates of sex (i.e. *σ* ≫ 1/*N*) on estimates of meiotic crossing over. (a) Decay of linkage disequilibrium over 900 sites, as a function of distance between two sites. Different colours denote individual rates of sex, as shown in the legend. Solid line is the mean value over 1,000 simulations; fainter curves represent 95% confidence intervals. 50 paired samples were simulated (100 samples in total), *N* = 10, 000, scaled mutation rate *θ* = 4*N_μ_* = 10, scaled crossover rate during sex *R* = 40. (b) As (a) but instead shows results from obligate sex simulations ran using ms, using a crossover rate equal to 40*σ* as shown in the legend. Due to binning of samples, *r*^2^ is shown for distances between 25 to 875 sites apart in (a) and (b). (c) Estimates of *R* using *LDhat*, as a function of the ‘effective’ crossover rate used in the facultative-sex coalescent simulation. Points are mean estimates from 1,000 simulations, bars are 95% confidence intervals. Black line denotes *y* = *x*; dashed red line is the linear regression fit. (d) Plot of all simulation results in (a) but instead as a function of the rescaled recombination rate 4*Ncσ* (plotted on a natural log scale), and after omitting polymorphisms with minor allele frequency less than 10%. Dotted lines show analytical expectations (Equation 6). Note the different y-axis scale compared to panel (a).

#### Low frequencies of sex

When the probability of sex is low (i.e., 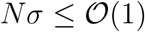), samples will diverge within individuals (Balloux *et al*. 2003; Bengtsson 2003; Ceplitis 2003; Hartfield *et al*. 2016). We examined how this allelic sequence divergence affects linkage disequilibrium by running simulations with 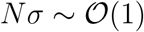 (specifically we investigated 2*Nσ* = Ω = 20, 2 and 0.2), but with a fixed scaled crossover rate 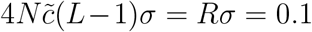. Although this scaled crossover rate is low, there is a high crossover rate when sex does occur, hence we expect to see some breakdown of linkage disequilibrium along the simulated genotype. We ran simulations over a larger number of sites (*L* = 100, 001) so there was enough distance to observe a decay in disequilibrium.

Figure 5(a) displays the linkage disequilibrium observed in low-sex cases, along with analytical expectations (Equation 7). After removing sites with minor allele frequency less than 10%, the Ω = 20 simulation exhibits *r*^2^ close to 0.45, which is as given by Equation 6 as the crossover rate goes to zero. However, lower rates of sex result in higher values of linkage disequilibrium, indicating that classic estimates of *r*^2^ using the rescaled recombination rate *Rσ* do not properly quantify disequilibrium when sex is low. Equation 7 captures the general behaviour of *r*^2^ under low frequencies of sex (i.e., elevated *r*^2^ values and a weaker dependence on the meiotic crossover rate) but there are several reasons why the results do not quantitatively match. Specifically, the analytical result is based on 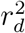 rather than *r*^2^, and finite sample sizes also introduce additional complications ignored in the calculation of expected 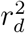. Our analytical model is intended to allow comparisons of the main patterns with the comparable sexual model, rather than providing precise predictions of the quantity as estimated by empiricists (which can instead be calculated using the simulation).

**Figure 5:**
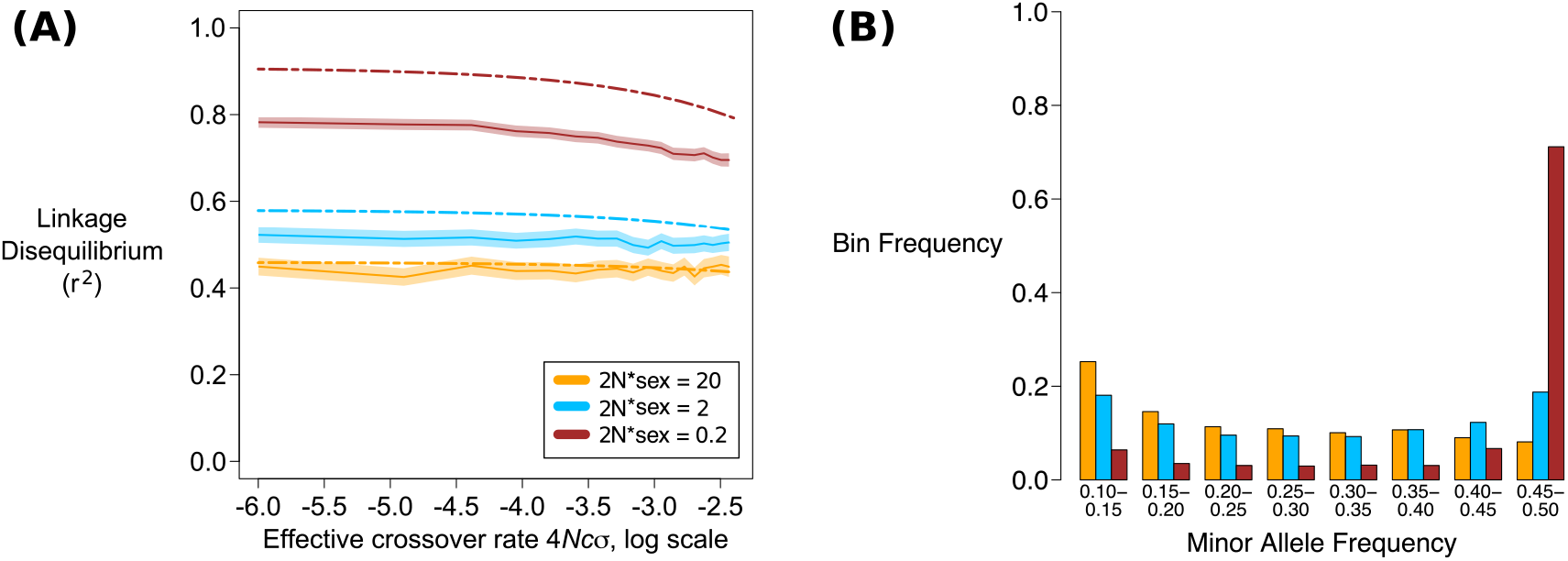
(a) Decay of linkage disequilibrium over 90,000 sites, as a function of the rescaled recombination rate 4*Ncσ* (on a natural log scale), and after omitting polymorphisms with minor allele frequency less than 10%. Different colours denote individual rates of sex, as shown in the legend. Solid line is the mean value over 1,000 simulations; fainter curves represent 95% confidence intervals. 50 paired samples were simulated (100 samples in total), *N* = 10, 000, scaled mutation rate *θ* = 4*N_μ_* = 10, scaled crossover rate over all 100,001 sites *Rσ* = 0.1. Coloured dash-dotted lines are low-sex analytical expectations for 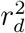 (Equation 7). *r*^2^ is shown for distances between 2,500 to 87,500 sites apart. Results for short distances (125 to 4,375 sites apart) are shown in Figure E in Supplementary File S2. (b) Histogram of minor allele frequencies for the low-sex scenarios; the bin frequency is measured over all 1,000 simulations. Bar colours correspond to the same rates of sex as used in panel (a).

Elevated *r*^2^ occurs under low sex due to allelic sequence divergence creating highly-differentiated haplotypes consisting of polymorphisms at intermediate frequencies (~50%). These polymorphisms arise due to a lack of genetic segregation creating highly differentiated haplotypes (Balloux *et al*. 2003; Bengtsson 2003; Ceplitis 2003; Hartfield *et al*. 2016). Figure 5(b) shows the density of minor allele frequencies over all simulation data, demonstrating that the Ω = 0.2 case has many sites with minor allele frequency between 45 and 50%. Consequently, *r*^2^ is higher over the genomic sample than expected based on obligate sex results using the effective crossover probability *cσ*. We also observe that linkage disequilibrium decay is only weakly affected by the meiotic crossover frequency, in line with analytical expectations (Equation 7).

### Linkage disequilibrium with mitotic gene conversion

#### High frequencies of sex

We ran simulations with mitotic gene conversion to investigate its effect on linkage disequilibrium. We define gene conversion using the population-level rate per sample, Γ = 4*N_g_*(*L* − 1). We first ran simulations with no meiotic crossovers (i.e., some degree of sexual reproduction occurs, but not any meiosis-related processes); here, the decay of linkage disequilibrium is independent of the rate of sex (provided sex is not too low; Figure 6(a)). This decay is similar to that observed in obligate sexual populations experiencing the same gene conversion rate (Figure 6(b)). When meiotic crossovers are included with rate *R* = 40, disequilibrium profiles separate out depending on the frequency of sex (Figure 6(c)), and are similar to those arising in obligate sexuals that experience the same gene conversion rate and an effective crossover probability *c_eff_* = *cσ* (Figure 6(d)). The pattern of linkage disequilibrium decay is more dependent on the probability of sex when the frequency of mitotic gene conversion is low, relative to the crossover probability *c* (contrast Figure 6 that uses Γ = 20, with Figure F in Supplementary File S2 that uses Γ = 2).

**Figure 6:**
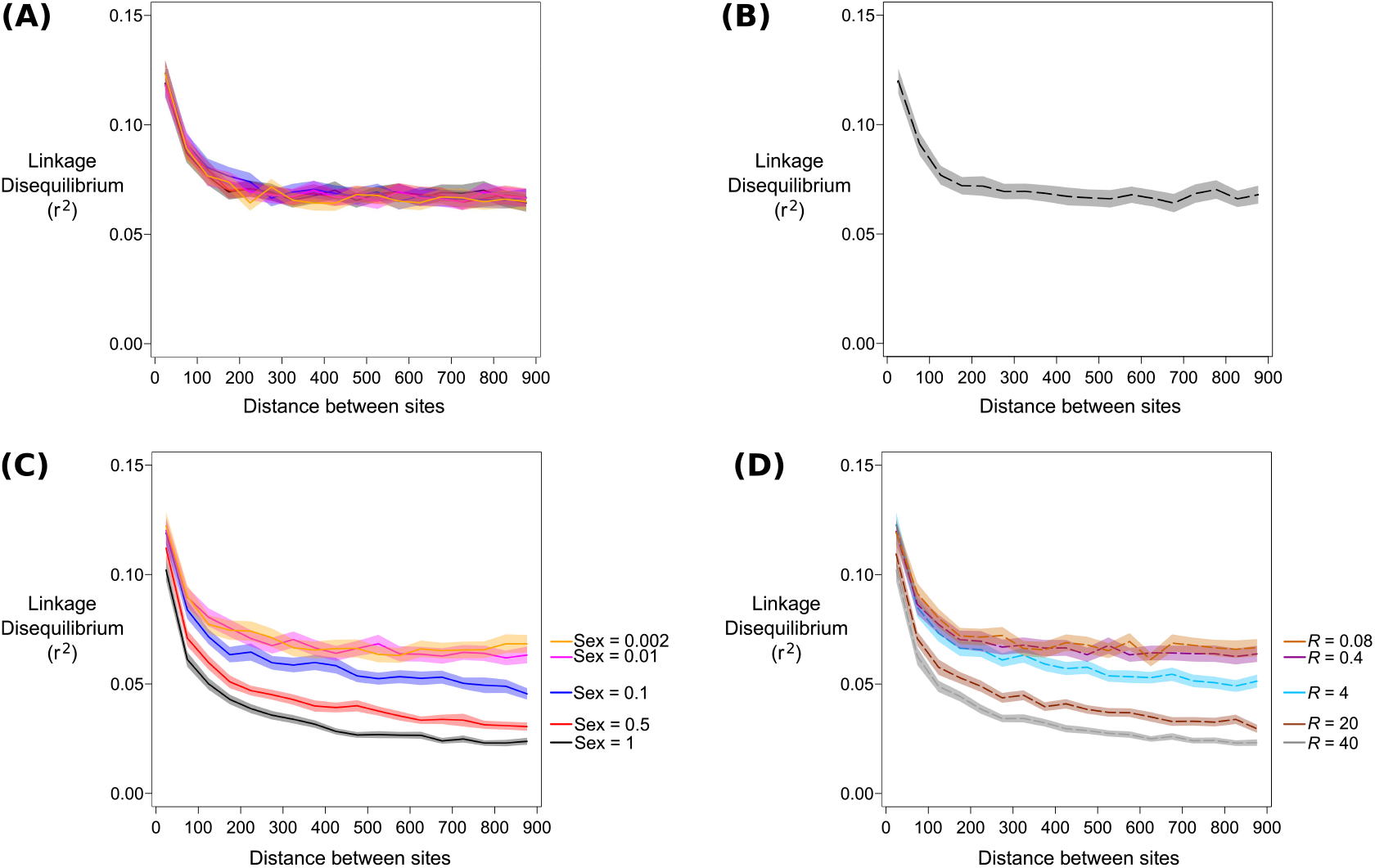
Decay of linkage disequilibrium over 900 sites, as a function of distance between two sites as caused by mitotic gene conversion with high rates of sex (*σ* ≫ *1/N*). 50 paired samples are taken from a population of size *N* = 10, 000, scaled mutation rate *θ* = 10, and mitotic gene conversion occurs with rate Γ = 20 (with average gene conversion tract length λ = 100 sites). Meiotic crossovers are either (a, b) absent or (c, d) present at rate *R* = 40. (a, c) Results from the facultative-sex coalescent simulation with different probabilities of sex. Colours are as shown in the legend; shaded bands are 95% confidence intervals. (b, d) Results from `ms` with 100 samples and the same gene conversion rate, with crossover probability *c_eff_* = *cσ* in (d). Note only one `ms` comparison is plotted in (b). Equivalent results with Γ = 2 are shown in Figure F in Supplementary File S2. *r*^2^ is shown for distances between 25 to 875 sites apart.

#### Low frequencies of sex

With low frequencies of sex 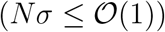, within-individual diversity is affected by the ratio of sex to gene conversion at a site, denoted *ϕ* (Hartfield *et al*. (2016); *ϕ* is defined mathematically below). If sex occurs more frequently than gene conversion (*ϕ* > 1), elevated within-individual diversity should be observed. However, if gene conversion arises at the same frequency, or more often than sex (*ϕ* ≤ 1), then gene conversion will lead to reduced within-individual diversity compared to a sexual population (Hartfield *et al*. 2016, Eq. 11). Hence we next ran simulations with different *ϕ* values to explore the relative effects of both phenomena on linkage disequilibrium.

We considered a diploid population *N* = 10, 000 from which we simulated 50 paired samples; *θ* = 10; and a genetic segment that is *L* = 10,001 sites long. To focus on the effects of gene conversion, we assumed no meiotic crossing over and only mitotic gene conversion was considered, with events having a mean tract length of λ = 1, 000 sites, matching estimates of non-crossover events obtained from yeast (Judd and Petes 1988; Martini *et al*. 2011). We fixed Ω = 2*Nσ* = 2 and varied Γ so that the ratio *ϕ* = (Ω*Q*)/Γ (for *Q* = (*L* − 1)/λ the number of breakpoints in units of mean gene conversion length), which determines neutral diversity at a single site, equals either 10, 1 or 0.1 (requiring Γ = 2, 20 and 200 respectively). Note that we define the probability of gene conversion per haplotype rather than per diploid genotype, so the probability of gene conversion is scaled by 4*N* as opposed to the 2*N* scaling used in Hartfield *et al*. (2016) (i.e., there is an extra factor of 2 in the denominator of *ϕ* to account for two haplotypes per individual).

Figure 7(a) demonstrates the unusual behaviour associated with high gene conversion with low rates of sex, with *r*^2^ being a non-monotonic function of the gene conversion frequency, in line with analytical findings (Figure 3(c)). Mitotic crossing over at rate 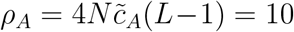 breaks down linkage disequilibrium over longer distance for Γ = 2 and 20, but not for Γ = 200 (Figure 7(b)). Both results are in line with analytical findings: *r*^2^ is a non-monotonic function of gene conversion when sex is rare (Figure 3(c)), and the presence of mitotic recombination can reduce long-distance *r*^2^, unless mitotic gene conversion acts at a much higher rate (Figure 3(d)).

**Figure 7:**
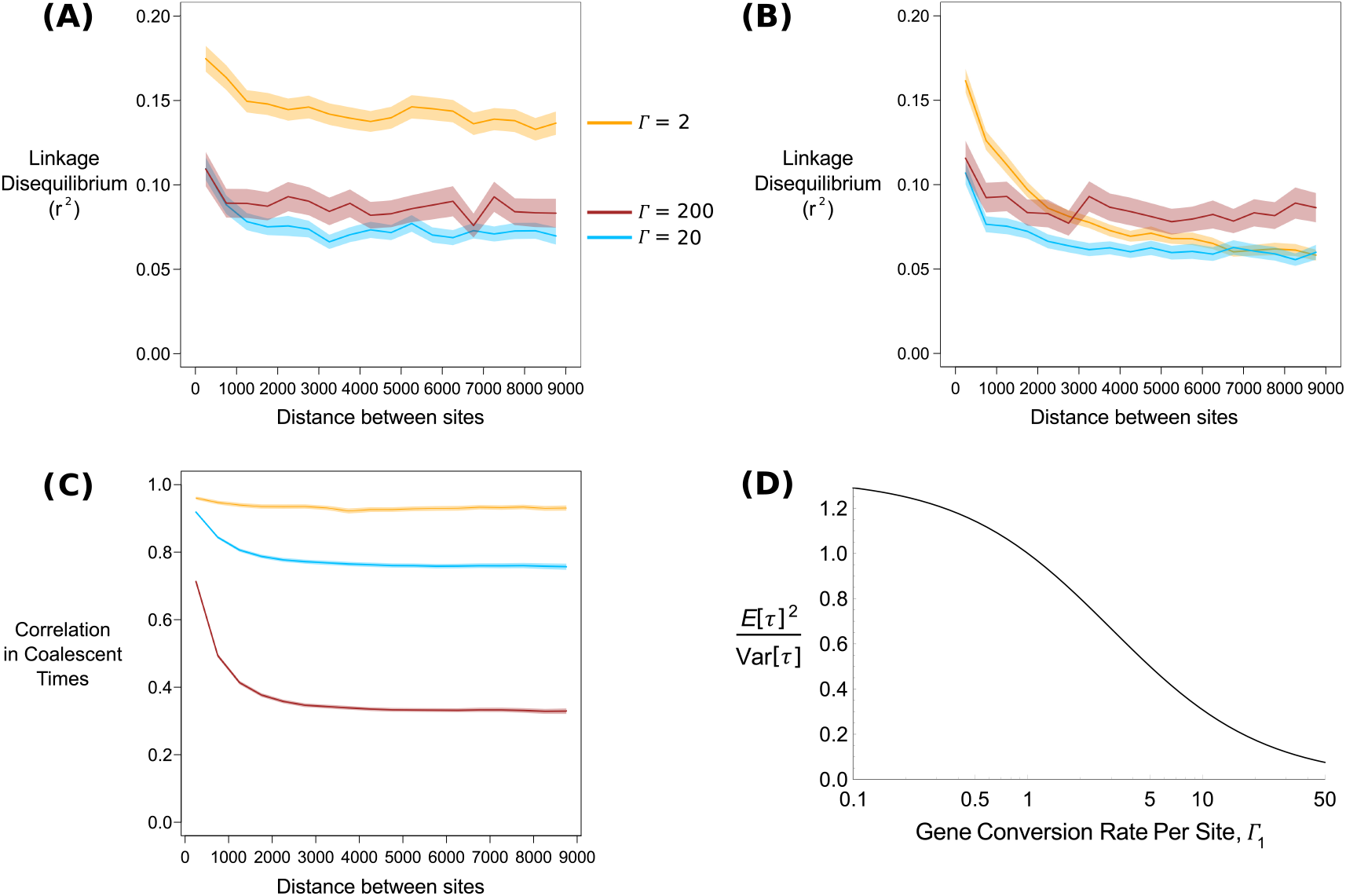
(a) Plot of linkage disequilibrium, measured using *r*^2^, as a function of distance between two sites. For a fixed rate of sex Ω = 2, gene conversion is set to Γ = 2 (orange line); 20 (blue line); or 200 (red line) with λ = 1, 000. Shading around lines indicate 95% confidence intervals. (b) As (a) but also including mitotic recombination with rate *ρ_A_* = 4*N*(*c_A_*) = 10. Results over short distances for (a) and (b) (25 to 875 sites apart) are presented in Figure G in Supplementary File S2. (c) Correlation in coalescent times (*Corr*[*ij,ij*] in Equation 8) between sites for the three Γ values, assuming no mitotic crossing over (i.e., *ρ_A_* = 0). Note that for Γ = 20 and 200, confidence intervals are only slightly thicker than the mean line. (d) Ratio of (*E*[*τ*]^2^/*Var*[*τ*]) for two samples taken from the same individual, as a function of the scaled gene conversion rate per site Γ_1_, with Ω = 2. (a)-(c) are shown for distances between 250 to 8,750 sites apart.

Elevated linkage disequilibrium is likely related to the reduced mean coalescence times that arise under frequent gene conversion. To further understand this behaviour, we can relate the observed *r*^2^ values to Equation 11 of McVean (2002), which demonstrated how 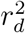 can be written as a function of both the correlation in coalescent times between sites, and the ratio of the mean coalescent time to the variance:

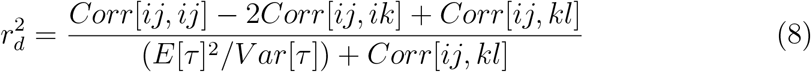

*Corr* in Equation 8 represent correlation in coalescent times between pairs of sites (e.g., *Corr*[*ij, kl*] the correlation in coalescence times where site one is taken from haplotype *i* and *j*, and site two is taken from haplotype *k* and *l*). *E*[*τ*] and *Var*[*τ*] are the mean and variance of coalescent times. Equation 8 shows that 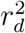 is not just reduced with lower covariances between pairs of loci, but it also decreases with higher *E*[*τ*]^2^/Var[*τ*]. This ratio equals one under the standard coalescent, but low sex alters the mean and variance of coalescent times (Hartfield *et al*. 2016) which will also affect this ratio, and subsequently alter linkage disequilibrium values. Figure 7(c) plots the covariance in coalescent times over all simulations, for two sites sampled from a single individual. We see that they are consistently lower with higher rates of gene conversion, reflecting how genetic material is more frequently transferred between samples. We next looked at the ratio (*E*[*τ*]^2^/*Var*[*τ*]), which can be calculated from Equations 11 and 12 of Hartfield *et al*. (2016). We focussed on the within-individual coalescence times, as these are directly affected by within-individual mitotic gene conversion. This ratio is shown in Figure 7(d) for Ω = 2 as a function of the mitotic gene conversion rate for a single site, Γ_1_. As Γ_1_ increases, the ratio (*E*[*τ*]^2^/*Var*[*τ*]) decreases, leading to the observed increase in *r*^2^ (Figure 7(d)). This result suggests that high rates of within-individual gene conversion distorts underlying genealogies, so that observed linkage disequilibrium is higher than that expected based on the rate of gene exchange alone. In contrast, meiotic crossing over has no direct effect on this ratio.

In Supplementary File S2 we investigate how linkage disequilibrium is affected if we alter *g* and λ while fixing the product *g*λ. Linkage disequilibrium decays more rapidly for higher *g* values with lower *λ* as there are more gene conversion events that break apart coalescent histories between individual sites.

### Effect of Population Subdivision

Measurements of linkage disequilibrium are known to increase under population structure with obligate sex (Wakeley and Lessard 2003), as polymorphisms that only appear in specific regions will naturally be in disequilibrium, increasing *r*^2^. Facultatively sexual organisms are known to show strong geographic differentiation (Arnaud-Haond *et al*. 2007). Hence we examined the effects of population structure in facultatively sexual organisms. We assumed an island model, consisting of 4 demes with a scaled migration rate *M* = 2*N_T_m* between them (for *N_T_* = 10, 000 the total population size across all demes). 50 paired samples were simulated, with 13 samples taken from two demes, and 12 from the other two. Population-scaled parameters are subsequently defined relative to *N_T_* (i.e., 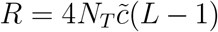, Ω = 2*N_T_σ*, Γ = 4*N_T_g*(*L* − 1)).

For high sex cases (*σ* ≫ 1/*N_T_*), and low sex cases 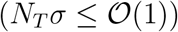 where mitotic gene conversion is present, results are qualitatively similar to those observed for a single population (Supplementary File S2). For the low-sex case with meiotic crossing-over, we ran simulations with *Rσ* = 0.1 and Ω equal to 20, 2 or 0.2, and compared them to an obligate sex case with the same crossover rate with different rates of migration. With high migration (*M* = 10) the results are similar to what is observed without population structure, with disequilibrium visually decaying along the genome sample for Ω = 0.2. Yet values are lower than in the panmictic case (compare the red line in Figure 8(a) with Figure 5). With lower migration (*M* = 0.1), disequilibrium values are unexpectedly reduced as the probability of sex decreases (Figure 8(b)). The reason for this unintuitive result is due to the partitioning of low-frequency polymorphisms under both low sex and population structure. With low migration rates, strong population structure is present so polymorphisms are localised to specific demes. Low frequencies of sex further partition polymorphisms within demes on diverged haplotypes (Figure 5(e) in Hartfield *et al*. (2016)). Hence the presence of rare sex, alongside high population structure, creates more polymorphisms at lower frequencies compared to populations with higher probabilities of sex (Figure 8(c, d)). These polymorphisms tend to have small values for *r*^2^, thereby reducing the average value. After removing polymorphisms with minor allele frequency less than 15%, estimates of *r*^2^ become similar for all rates of sex, although Ω = 0.2 results are still slightly lower than other cases for *M* = 0.1 (Figure J in Supplementary File S2).

**Figure 8:**
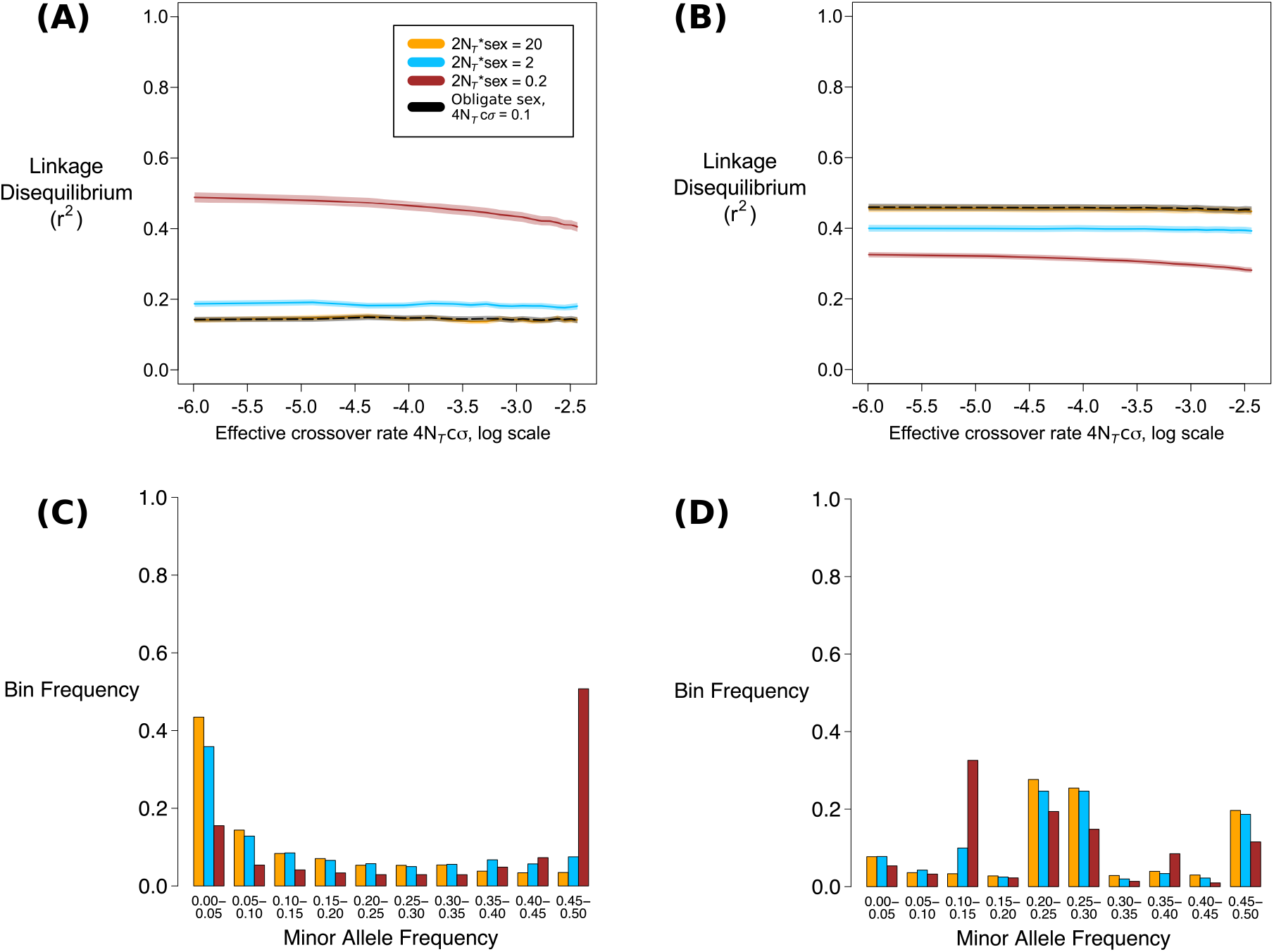
(a, b) Decay of linkage disequilibrium over 90,000 sites from samples taken over a subdivided population, as a function of the rescaled recombination rate 4*N_T_cσ* (plotted on a natural log scale). Different colours denote individual rates of sex, as shown in the legend. Solid line is the mean value over 1,000 simulations; fainter curves represent 95% confidence intervals. 50 paired samples were simulated (100 samples in total) over 4 demes, *N_T_* = 10, 000, scaled mutation rate *θ* = 4*N_T_μ* = 10, scaled crossover rate over entire ancestral tract *Rσ* = 0.1, scaled migration rate is either (a) *M* = 10 or (b) *M* = 0.1. Black dashed line is equivalent obligate sex simulation ran using `ms` with 100 samples. *r*^2^ is shown for distances between 2,500 to 87,500 sites apart. (c, d) Histogram of minor allele frequencies, with the bin frequency measured over all 1,000 simulations. Bar colours correspond to the same rates of sex as used in panels (a, b).

## Discussion

### Summary of results

Existing single-locus theory for facultatively sexual organisms shows behaviour distinct from sexual cases only with extremely low frequencies of sex 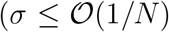; Bengtsson (2003); Ceplitis (2003); Hartfield *et al*. (2016)). In this paper we provide novel analytical and simulation results to investigate how correlations in genetic diversity across loci are affected by facultative sex. We also provide an updated version of a simulation package, and explain how existing crossover (Hudson 1983) and multi-site gene conversion routines (Wiuf and Hein 2000) can be included in facultative-sex coalescent processes, to investigate how they affect gene genealogies. This program can be used to simulate ancestral recombination graphs of facultatively sexual organisms.

When the frequency of sex is high (*Nσ* ≫ 1), we observe that the breakdown in linkage disequilibrium in a genetic sample is similar to that observed in an obligate sex model using an effective crossover probability *c_eff_* = *cσ* (Figure 3(a), Figure 4). This result reflects similar behaviour in partially self-fertilising organisms (Nordborg 2000), where the effective crossover rate is equal to *r*(1 − *F*) for *F* the inbreeding rate (although this scaling breaks down with high self-fertilisation and crossover rates; Padhukasahasram *et al*. (2008); Roze (2009, 2016)).

Hence if there exists knowledge of meiotic crossover rates, then one can use linkage disequilibrium data to estimate the overall frequency of sex. The situation becomes more complicated if mitotic recombination is present as it also breaks down linkage disequilibrium, even under low frequencies of sex. If sex is frequent but crossing over is rare, mitotic gene conversion principally affects linkage disequilibrium (Figure 6(a)). Once crossing-over probabilities become high then these principally break down linkage disequilibrium, so the effective crossover rate scaling *c_eff_* = *cσ* still holds (Figure 6(b)).

When rates of sex become low 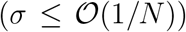, the decay in linkage disequilibrium can no longer be captured by rescaling *c_eff_* = *cσ*, as the distribution of genealogies becomes fundamentally different than when sex is common (Figure 3(b-d)). In the absence of gene conversion, *r*^2^ becomes elevated with low rates of sex, reflecting more linked polymorphisms present at intermediate frequencies (Figure 5). If mitotic gene conversion is present, the ratio between rates of sex and gene conversion *ϕ* becomes a strong determinant of linkage disequilibrium, with unexpected behaviour arising if gene conversion occurs at high rates relative to sex. Increasing gene conversion will first reduce overall disequilibrium values due to gene exchange breaking down associations between sites. Yet very high rates of gene conversion then cause elevated linkage disequilibrium values, which is a consequence of how gene conversion changes the distribution of coalescence times. Adding mitotic crossovers reduces the minimum observed linkage disequilibrium, unless mitotic gene conversion occurs at much higher rates (Figures 3(c, d) and 7). Finally, low sex combined with low migration rates in subdivided populations also reduces *r*^2^ values, due to more low-frequency polymorphisms being present within demes (Figure 8). These non-intuitive effects illustrate the value of explicitly modelling genetic diversity under low rates of sex when considering genomic data for facultatively sexual organisms.

### Future Directions

The creation of the new coalescent algorithm that accounts for facultative sex, crossing over and gene conversion can be used as a basis for inferring these processes from genomic data. This can be achieved by using coalescent simulations to create likelihood profiles over two loci (Hudson 2001; McVean *et al*. 2002; Wall 2004; Auton and McVean 2007). An alternative approach would be to use Approximate Bayesian Computation to recurrently simulate different outcomes, each time comparing them to the real data and keeping those that match sufficiently well, to build a pseudo-likelihood (Sunnåker *et al*. 2013). Simulation results also suggest that it is important to jointly consider both genome-wide diversity and linkage disequilibrium if we wish to infer the effects of sex, meiotic crossovers, and gene conversion, especially if mitotic gene conversion is pervasive (Figure 7).

We anticipate that the *FacSexCoalescent* simulation can be expanded upon in the future to account for more complex scenarios. The only population structure we considered is an island model, whereas bottlenecks or unequally-sized subpopulations are common (Pool and Aquadro 2006; Veeramah and Hammer 2014; Frantz *et al*. 2016). The gene conversion model can also be expanded upon to consider context-dependent events (for example, GC-biased gene conversion; Duret and Galtier (2009)). Given ongoing debates on how gene conversion potentially affects genetic diversity and fitness in facultatively sexual organisms (Mancera *et al*. 2008; Flot *et al*. 2013; Tucker *et al*. 2013), a deeper understanding of how gene conversion affects the distribution of genetic diversity can shed further insight into what processes influence genetic evolution in facultatively sexual organisms.

## Acknowledgements

We would like to thank two anonymous reviewers and Associate Editor John Wakeley for providing constructive comments on the manuscript. MH was supported by a Marie Curie International Outgoing Fellowship, grant number MC-IOF-622936 (project SEXSEL), and an European Research Council grant (FP7/20072013, ERC Grant 311341) awarded to Thomas Bataillon. This work was also supported by Discovery Grants (AFA & SIW) from the Natural Sciences and Engineering Research Council of Canada.

## A Implementing Recombination into the Facultative Sex Simulation Algorithm

### Overview of basic coalescent simulation

Here we outline the implementation of meiotic and mitotic recombination events in the facultative sex coalescent simulation routine (Hartfield *et al*. 2016). We describe the probability that set events occur per generation; that is, both the time in the past to the next event and resolution of events are based on unscaled probabilities (as opposed to rates, where a probability is multiplied by the population size to give the expected number of events per generation). We define *p_NS_* as the probability that none of the *x* paired samples are split by sex and *p_E0_* as the probability of any event (e.g., coalescence, recombination) given that none of the paired samples are affected by sex. The absolute time to the next event is drawn from a geometric distribution with parameter *p_sum_* = (1 − *p_NS_*) + *p_NS_p_E0_*; this time is rescaled by 2*N* so that it is on a coalescent timescale. It is subsequently determined whether any and, if so, how many of the *x* paired samples segregate into different individuals due to sexual reproduction. If *k* of *x* paired samples are produced via sex, then 2*k* new unpaired samples are created. The total probability of any other event occurring is then re-calculated, conditional on this updated sample configuration. It is determined whether any such event occurs, and which type if one does arise; the sample configuration is updated appropriately. Note that if sex is common, the first term in *p_sum_* is large and all paired samples are rapidly split by sex, so the model then behaves like a haploid process. If the population is structured as an island model, the logic is similar but we instead track *x_i_, y_i_* paired and unpaired haplotypes in deme *i*, and consider 2*N_T_* total haplotypes over all subpopulations. We refer the reader to Hartfield *et al*. (2016) for further details of the basic coalescent simulation.

### Implementing Meiotic and Mitotic Crossing Over

We outline the probability of either a crossover or gene conversion event occurring each generation, then implement these probabilities into the calculation of *p_sum_* as described above. As with the single-locus routines, we assume that sexual reproduction occurs first, followed by subsequent gene exchange events. Let 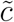 be the absolute meiotic crossover probability between any two adjacent sites, conditional on sex having occurred; 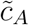 the mitotic crossover probability (which is not conditional on the reproductive mode); and *L* the number of sites that the genetic samples cover. Assuming 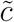 and 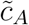 are small, the total meiotic crossover probability in each individual at the start of the process is 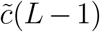. We assume that the total recombination probability is low (i.e., 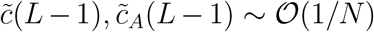) so we do not consider outcomes where more than one crossover event occurs per generation.

Following sexual reproduction, crossovers act on unpaired samples with probability 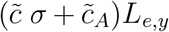. The quantity *L_e,y_* is the ‘effective’ crossover length summed over all *y* unpaired samples. We include this term to ensure that unnecessary crossover events are not considered, thus speeding up the routine (Hein *et al*. 2005). *L_ey_* is defined as follows: let *L_s,i_* be the first ancestral site in unpaired sample *i*, and *L_t,i_* the last ancestral site. Then *L_e,i_* = *L_t,i_* − *L*_*s,i*_ equals the total number of breakpoints where a crossover can create two new samples, each carrying ancestral material; note that any sites within individual haplotypes that have reached their most recent common ancestor are converted into non-ancestral material (Hein *et al*. 2005). Then *L_e,y_* = ∑_*i*∈*y*_ *L_e,i_*. This crossover event creates two new samples, with each part carrying ancestral material (Figure 1(a)).

If *k* out of the *x* paired samples segregate via sex into 2*k* new unpaired samples, then the crossover probability is increased by adding on an extra 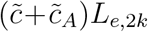 term. Here *L_e,2k_* is the ‘effective’ crossover length over the 2*k* new unpaired samples, defined in a similar manner to *L_e,y_*. The 2*k* samples are a transitory class of unpaired haplotypes, created through sexual reproduction segregating paired samples into distinct individuals (see also Figure 1(b, c)). Because they are already determined to be have been created by sex by an earlier step in the algorithm, there is no factor *σ* contributing to their probability of experiencing meiotic crossing over. Those that do not undergo crossing-over become regular unpaired samples (Figure 1(b)); those that do are transformed into regular paired samples (Figure 1(c)).

Mitotic crossing over can act on the remaining *x* − *k* paired samples that do not undergo sexual reproduction. This event occurs with probability 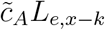, for *L_e,x−k_* the effective crossover length measured over both arms within the remaining *x* − *k* paired samples. *L_e,x−k_* is measured in a different manner than for unpaired samples. Let *i* be an individual where both haplotypes *i*_1_, *i*_2_ are sampled. Define *L_s,i_1__, L_s,i_2__* as the first ancestral site in each of these samples, and *L_t,i_1__, L_t,i_2__* the last ancestral sites. Then the first ancestral site at which mitotic crossing over is valid in individual *i* is *L_s,i_* = *min*(*L_s,i_1__, L_s,i_2__*); similarly, *L_t,i_* = *max*(*L_t,i_1__, L_t,i_2__*). Then *L_e,i_* = *L_t,i_* − *L_s,i_* and *L_e,x−k_* = ∑_*i∈(*x−k*)*_ *L_e,i_*. Mitotic crossing over exchanges genetic material between the two samples within an individual (Figure 1(d)).

These probabilities are considered alongside other events to determine whether the next event involves a meiotic crossover. If it is chosen, then one of the appropriate samples is picked at random (weighing by the length of extant breakpoints present in each sample), and the appropriate outcome is enacted. Note that if the potential for crossing over is high (i.e. if the probability of sex and crossing over is high, and there are a large number of samples) then the net recombination probability can exceed one, as the assumption that only up to one recombination event occurring per generation is violated, causing the algorithm to terminate. Hence large crossover probabilities should be avoided.

### Implementing Meiotic, Mitotic Gene Conversion

To account for both meiotic and mitotic gene conversion events (Figure 1(e-g)), up to four additional parameters are specified. Two new parameters are *gS* and *g*, the probabilities of either meiotic or mitotic gene conversion occurring with its leftmost boundary arising on the recipient homolog at a given site. We also define the average length of gene conversion events, denoted λ_*S*_ for meiotic gene conversion and λ for mitotic gene conversion. We implement and extend the algorithm of Wiuf and Hein (2000) to calculate the probability of either type of gene conversion event occurring each generation. Here, the length of gene conversion events (scaled by the total number of breakpoints) is drawn from an exponential distribution with parameter *Q* = (*L* − 1)/λ, the number of breakpoints in units of average gene conversion length (Wiuf and Hein 2000). We define distinct *Q_S_* = (*L* − 1)/λ_*S*_, *Q* = (*L* − 1)/λ for meiotic and mitotic gene conversion events. Further details of the mathematical derivations are in Section B of Supplementary Mathematica File S1.

There also exists a special class of gene conversion events, where conversion initiates outside the ancestral tract and extends completely over ancestral material (Figure 1(h)). If there exist *x* − *k* pairs after *k* of them are split by sex, then the probability of this event happening equals 2(*x* − *k*)(*g*(*L* − 1)*e*^−*Q*^)/*Q* (a full derivation is presented in “Deriving probability of ‘complete’ gene conversion” below, and Section C of Supplementary Mathematica File S1).

### Determining type of gene conversion (meiotic or mitotic)

To understand how the different events (meiotic and mitotic gene conversion) are considered, it is easiest to relate calculations to the obligate sex case with a single type of gene conversion event. Here, the product of the gene conversion probability *g*_0_ (using *g*_0_ to differentiate this general gene conversion probability from the mitotic gene conversion notation) and the number of breakpoints *L* − 1 is such that 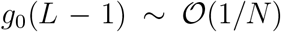. Then the probability of a disruptive gene conversion event in a sample of length *L* is 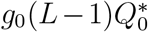, where *g*_0_ is the probability that the leftmost edge of a gene conversion tract is at a given site, and:

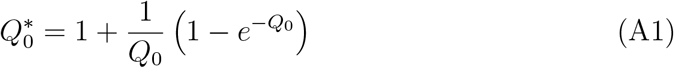

*Q*_0_ = (*L*−1)/λ_0_ is the number of breakpoints in units of average gene conversion length (here too, we use *Q*_0_, 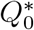 and λ_0_ to define this general gene conversion process). 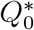 accounts for gene conversion events that only partly lie in ancestral material (i.e., only one end of the gene conversion lies in ancestral material) as well as those that lie entirely within this region (i.e., both breakpoints lie within ancestral material). Equation A1 assumes that the length of gene conversion events (scaled by the total number of breakpoints) is drawn from an exponential distribution with parameter *Q*_0_ (Wiuf and Hein 2000). Equation A1 also disregards possible edge effects (e.g., if the ancestral tract lies near the chromosome edge). In the facultative-sex coalescent, we can partition this probability depending on the type of conversion event (meiotic or mitotic), and the number of each type of sample (paired or unpaired) present at the time. Let there be (*x* − *k*) paired samples after *k* of them have split after sex, making 2(*x* − *k*) haplotypes in total; *y* unpaired samples; and 2*k* new unpaired samples following genetic segregation. After partitioning over all possible outcomes, the total probability of disruptive gene conversion equals:

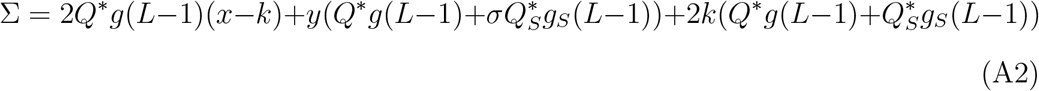

Here, *Q*\* and 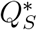 are equal to 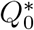 above, with parameters *Q* = (*L* − 1)/λ and *Q_S_* = (*L* − 1)/λ_*S*_ for mitotic and meiotic gene conversion respectively. As segregation has already been resolved, the (*x* − *k*) remaining paired samples reproduce asexually so only they can be subject to mitotic gene conversion. Unpaired samples can be subject to both meiotic and mitotic gene conversion, hence for each unpaired sample (*y* in total) we also have to consider the probability of sex *σ*. Note that there is no *σ* term when considering the 2*k* new unpaired samples, as they have already undergone sex by this point in the algorithm. In contrast to the crossover procedure, we do not weigh samples by the number of breakpoints within ancestral material; gene conversion events affecting only non-ancestral material are allowed to occur.

When a disruptive gene conversion event occurs, it is first determined if it acts on unpaired or paired samples. The probability that gene conversion acts on a paired sample is 2*Q*g*(*L* − 1)(*x* − *k*)/Σ where Σ is given by Equation A2, and one minus this probability is the chance it acts on unpaired samples. If acting on a paired sample then only mitotic gene conversion can occur. If acting on unpaired samples, a further random draw is made to determine whether the gene conversion event is meiotic or mitotic. Let 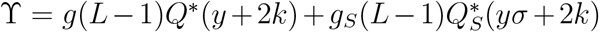 be the probability of a gene conversion event that occurs on an unpaired sample. If an unpaired sample undergoes conversion, the probability that the event is mitotic equals (*g*(*L* − 1)*Q*\*(*y* + 2*k*))/ϒ; a similar calculation can be made for meiotic gene conversion.

### Drawing start, end breakpoints following gene conversion

The scaling terms *Q*\*, 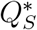 account for the fact that gene conversion does not necessarily take place entirely within the gene tract, but may only partially overlap with it. We follow the logic outlined in Wiuf and Hein (2000) to accurately model the relative frequency of each of these events. Given the tract length in units of conversion events *Q*_0_ (which can be either *Q* or *Q_S_*), *K*(*Q*_0_) = 1 − (1 − exp(*Q*_0_))/*Q*_0_ is the probability that if gene conversion starts in the sample, it will also end within it (Wiuf and Hein 2000, Eq. 2). One can then define the probability that both breakpoints occur within the sample (Wiuf and Hein 2000, Eq. 11):

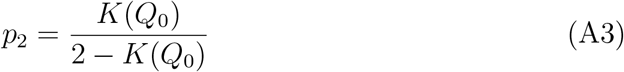

The probability that only one breakpoint falls within the sample equals *p*_1_ = 1 − *p*_2_ (Wiuf and Hein 2000, Eq. 12). We first choose whether one or both breakpoints lie within the sample, as determined by Equation A3. The appropriate start and end points are then chosen from the relevant probability distributions. Wiuf and Hein (2000) determined how the distribution of breakpoints depends on whether one or both breakpoints lie within the genome tract. For example, if only one breakpoint lies in the tract then it is likelier to occur closer to one of its edges. When choosing gene conversion breakpoints, they are selected by calculating the cumulative distribution function (CDF) for the event; drawing an initial start or end point from a uniform distribution; then plugging this uniform draw into the inverse CDF to obtain the true start or end point. The CDFs are obtained from the relevant probability distribution functions outlined by Wiuf and Hein (2000). Note that the resulting outputs are continuous variables lying between 0 and 1, while the simulation program assumes discrete breakpoints. Hence after the relevant breakpoint locations have been found, it is then converted into the appropriate discrete value lying between 1 and *L* − 1, including these values. Further details on the following derivations are provided in Section B of Supplementary Mathematica File S1.

If *two breakpoints are chosen*, then the joint probability distribution of start points *s* and end points *t* equals *f*(*s,t*) = (*Q*_0_ exp(−*Q*_0_(*t* − *s*)))/*K*(*Q*_0_) (Wiuf and Hein 2000, Eq. 4). By integrating out *t* from *s* to 1, one obtains the marginal density of start points, *f*(*s*) = (1 − exp(−*Q*_0_(1 − *s*)))/*K*(*Q*_0_) (Wiuf and Hein 2000, caption of Fig. 4). The CDF of *s* can then be calculated as:

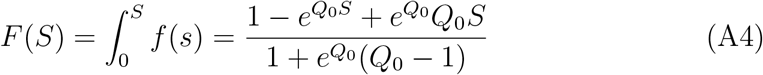

To choose a start point, we draw a value between 0 and 1 from a uniform distribution and plug it into *F*^−1^(*S*), which equals:

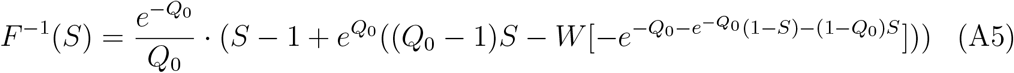

where *W* is the Lambert function (Abramowitz and Stegun 1970). To draw the respective end point, we first need to determine the distribution *f*(*t|s*); i.e., the density of end points *given* a starting point *s*. This function is also equal to *f*(*s,t*)/*f*(*s*) = (*Q*_0_exp(−*Q*_0_(*t* − *s*)))/(1 − exp(−*Q*_0_(1 − *s*))). From this function we obtain the CDF of *T* given *s*, as well as the inverse CDF:

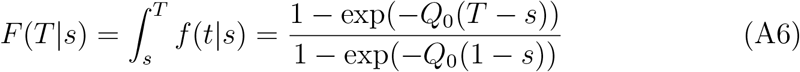

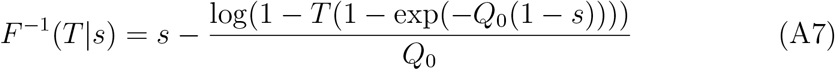

Equation A7 is then used to determine the endpoint of of the gene conversion event, which automatically lies within the length of the genetic sample. If the chosen endpoint is the same as the startpoint, then another endpoint is chosen so that they are distinct.

If *one breakpoint is chosen*, it can be the start or end of gene conversion with equal probability (Wiuf and Hein 2000, Eq. 7). If it is chosen to be the end point of gene conversion initiating outside the tract, then the start point is set to zero (i.e., the far left edge of the tract). The probability density of end points *t* is *f*(*t*) = exp(−*Q*_0_*t*)/(1 − *K*(*Q*_0_)) (Wiuf and Hein 2000, Eq. 8). This function is left-skewed; end points are likely to be near the left-hand side of the genetic sample. The CDF and inverse CDF can be calculated as:

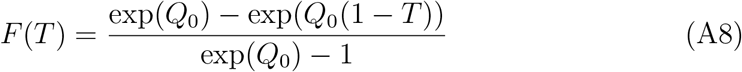

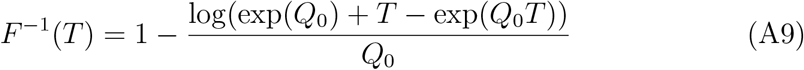

If the single breakpoint is instead a start point, then the end point is set to the extreme right side of the sample. The probability density of start points is given by *f*(*s*_1_) = exp(−*Q*_0_(1 − *s*_1_))/(1 − *K*(*Q*_0_)) (Wiuf and Hein 2000, Eq. 6). This function is right-skewed; start points are likely to appear towards the end of the sampled genome. The CDF and inverse CDF equal:

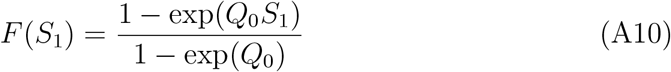

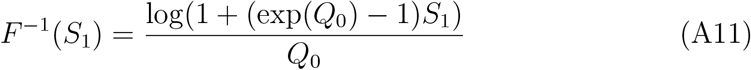

Before gene conversion is carried through, it is first checked whether it would result in a sample that does not carry any ancestral material. These fully non-ancestral samples can arise if either (i) conversion acts on an unpaired sample, in a region spanning entirely non-ancestral material; or (ii) conversion acts over all remaining ancestral material in a paired sample, rendering it non-ancestral. In case (i) the action stops without creating this ‘ghost’ sample. In case (ii) gene conversion causes a within-individual coalescent event, converting the paired sample into an unpaired sample. The recipient sample becomes non-ancestral and is no longer tracked.

### Deriving probability of ‘complete’ gene conversion

Let *Q* be the mean scaled length of mitotic gene conversion events. Following Wiuf and Hein (2000), the length of gene conversion events can be drawn from an exponential distribution with parameter *Q*. Let a gene conversion event start at a distance *x* from the focal sequence (where distances are scaled by the number of breakpoints *L* − 1). The gene conversion event will therefore cover the focal sequence with probability *e*^−*Q*(1+*x*)^. The probability of a complete gene conversion occurring over all paired haplotypes (of which there exist 2(*x* − *k*)), and over the entire density of conversion breakpoints (of which there exists *g*(*L* − 1) per length of focal sequence) equals 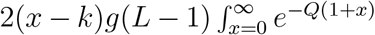. Solving the integral gives the probability 2(*x* − *k*)(*g*(*L* − 1)*e*^−*Q*^)/*Q*.

Note that if *L* > 1 then this probability goes to infinity as *Q* → 0. In this case, the average gene conversion length is much larger than the genetic sample being simulated (i.e., λ ≫ *L* − 1), so any gene conversion event is likely to affect the entire genetic region. The reason this discontinuity arises is because the coalescent process assumes that no more than one event occurs per generation. Wiuf and Hein (2000) ensures this logic is maintained by assuming 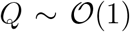. Furthermore, a small *Q* value would invalidate the assumption used to compute *Q*\*; specifically, conversion events that initiate outside the sample but end within it only do so in regions near to the genetic sample (since the probability of these events equals *e*^−*nQ*^ if initiating *n*(*L* − 1) breakpoints away from the sample). Hence while the simulation can be run with very small *Q* values, it is inadvisable to do so as erroneous genealogies may be produced.

